# Importance of AB domain in parvalbumins’ calcium binding affinity

**DOI:** 10.1101/2022.05.27.493786

**Authors:** Kalyan Immadisetty, Jeremiah Jacob-Dolan

## Abstract

Members of the parvalbumin (PV) family of calcium binding proteins are found in a variety of vertebrates, where can they influence neural functions, muscle contraction and immune responses. It was reported that the *α*-parvalbumin (*α*PV)s AB domain comprising two *α*-helices, dramatically increases the proteins calcium (Ca^2+^) affinity by ≈10 kcal/mol. To understand the structural basis of this effect, we conducted all-atom molecular dynamics (MD) simulations of WT *α*PV and truncated *α*-parvalbumin (Δ*α*PV) constructs. Additionally, we also examined the binding of magnesium (Mg^2+^) to these isoforms, which is much weaker than Ca^2+^ (Mg^2+^ actually does not bind to the Δ*α*PV). Our key finding is that ‘reorganization energies (RE)’ assessed using molecular mechanics generalized Born approximation (MM/GBSA) correctly rank-order the variants according to their published Ca^2+^ and Mg^2+^ affinities. The 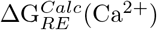 of the Δ*α*PV compared to the wild-type (WT) is 415.57±0.55 kcal/mol, indicating that forming a holo state of Δ*α*PV in the presence of Ca^2+^ incurs a greater reorganization penalty than the WT. This is consistent with the Δ*α*PV exhibiting lesser Ca^2+^ affinity than the WT (≈9.5 kcal/mol). Similar trend was observed for Mg^2+^ bound variants as well. Further, we screened for metrics such as oxygen coordination of EF hand residues with ions and found that the total oxygen coordination number (16 vs. 12 in WT:Ca^2+^ and Δ*α*PV:Ca^2+^) correlate with the reported ion affinities (−22 vs. −12.6 kcal/mol in WT:Ca^2+^ and Δ*α*PV:Ca^2+^), which indicates that AB domain is required for the protein to coordinate with maximal efficiency with the binding ions. To our surprise, no significant differences were observed between the Mg^2+^ bound WT and Δ*α*PV isoforms. Additionally, we have screened for factors such as total number of waters, hydrogen bonds, protein helicity and *β*-content for the entire protein, which enables us to understand the impact of lack of AB domain on the entire structure and not just binding sites. Our data indicate that AB improves the overall helicity (≈5%) in apo as well as holo forms. Particularly, AB increases *α*-helicity in the D-helix residues (i.e., 60–65) upon ion binding by ≈35% (90% vs. 55% in the Ca^2+^ bound WT and Δ*α*PV, 60% vs. 20% in the Mg^2+^ bound WT and Δ*α*PV), which likely contributes to high Ca^2+^ binding affinity. On the contrary, no significant effect on the overall *β*-content was observed. Similarly, increased dehydration (≈50) and increase in total number of hydrogen bonds (≈7) were observed upon ion binding in both the WT and Δ*α*PV systems, however, no significant differences were observed between the WT and Δ*α*PV variants and also between Ca^2+^ and Mg^2+^ isoforms. We speculate that this is due to the partially folded apo state that was captured in our MD simulations, which might not be physiologically relevant as suggested by NMR experiments [1]. Also, we have identified seven different interactions that might play a key role in binding the AB domain with the CDEF helices, particularly the D22(AB)–S78(CDEF) hydrogen bond. Overall, this study indicates that local (i.e., the EF hands) as well as global factors play a role in improved ion binding due to AB domain.

## 2 Introduction

Parvalbumin (PV) is a Ca-binding protein (CBP) protein and shape Ca^2+^ signaling in many physiological processes such as neural function, muscle contraction, and immune responses [2]. PV isoforms display dramatically different Ca^2+^ and Mg^2+^ affinities [3, 4, 5, 6]. PVs are popular targets for gene therapy for diseases involving calcium dysregulation [7, 8] or as biochemical and structural characterization to elucidate the molecular basis of Ca^2+^-affinity [9, 10, 11, 12, 13, 14].

Alpha PV (*α*PV), a member of the PV family, is expressed in mammalian skeletal muscle [15], GABA-ergic neurons and the organ of Corti [16]. *α*PV was implicated in accelerating the relaxation of muscle contraction through binding calcium [17]. *α*PV selectively binds Ca^2+^ over Mg^2+^ by ≈10 kcal/mol [18, 19], and is uniquely a strong binder of Ca^2+^ compared to other PV isoforms. The intrinsic ability of *α*PV to accelerate muscle relaxation has motivated efforts to engineer and target engineered variants to cardiac tissue as a therapy for diastolic dysfunction [8], or for detailed mutagenesis studies [19, 18] to elucidate its Ca^2+^ binding properties.

Predicting the energetics of Ca^2+^ and Mg^2+^ binding and the structural changes that follow has remained challenging. Bioinformatics techniques facilitated the identification of ion binding domains based on protein sequences [20, 21, 20]. Algorithms accounting for the geometric configuration of Ca^2+^ chelating ligands to achieve the optimal coordination geometry have further improved EF-hand prediction [22], and helped rationalizing the selectivity among other ions such as Mg^2+^ [23]. MD simulations [24] have facilitated detailed analyses of Ca^2+^ binding in intact proteins. More recent applications include refinement of force field parameters for Ca^2+^ [25, 26], Ca^2+^ binding energetics in EF-hand containing proteins [27, 28, 29, 30], effects of Ca^2+^ binding on protein function [31, 32, 33], mutations that disrupt the binding of Ca^2+^ [34, 30] and differential modulation of purinergic receptorsn by Mg^2+^ and potassium (K^+^) [35].

In this study, we applied MD simulations and free energy calculations to rationalize the differences in Ca^2+^ and Mg^2+^ binding between wild type *α*PV and Δ*α*PV. Δ*α*PV is a truncated variant of *α*PV, which lacks two helices (A,B). The truncated construct (Δ*α*PV) exhibited dramatically reduced Ca^2+^ binding [1] and do not bind to the Mg^2+^. This study indicates that the two helices (A,B) strongly contribute to Ca^2+^ binding and selectivity in *α*PV.

## 3 Results

The goal of this study was to identify structural factors that contribute to high Ca^2+^ binding affinities and selectivities versus Mg^2+^ in intact *α*PV and Δ*α*PV, and also the role of AB domain on the respective ion binding affinities. We performed ≈18 *μ*s of unbiased MD simulations for the Ca^2+^-bound, Mg^2+^-bound and apo states of these respective proteins. We then compared Δ*α*PV to the WT *α*PV via thermodynamic scores from MM/GBSA and compared against trends based on their experimentally determined Ca^2+^ and Mg^2+^ affinities. We next decompose these scores into contributions from the EF-hand ion coordination sites versus structural and energetic changes elsewhere in the protein to rationalize experimentally observed binding trends.

### 3.1 Comparing simulated *α*PV structures

We first verify that the MD predicted structures resemble those deposited in the Protein Databank. In Fig. 1A,B we demonstrate that the predicted holo (red) structures are generally consistent with their respective experimental reference structures (PDB:1RWY [36] and PDB:1G33 [1] (green)), except for the loss of helicity in the D-helix in the Δ*α*PV(purple colored circle in Fig. 1B).

**Figure 1:**
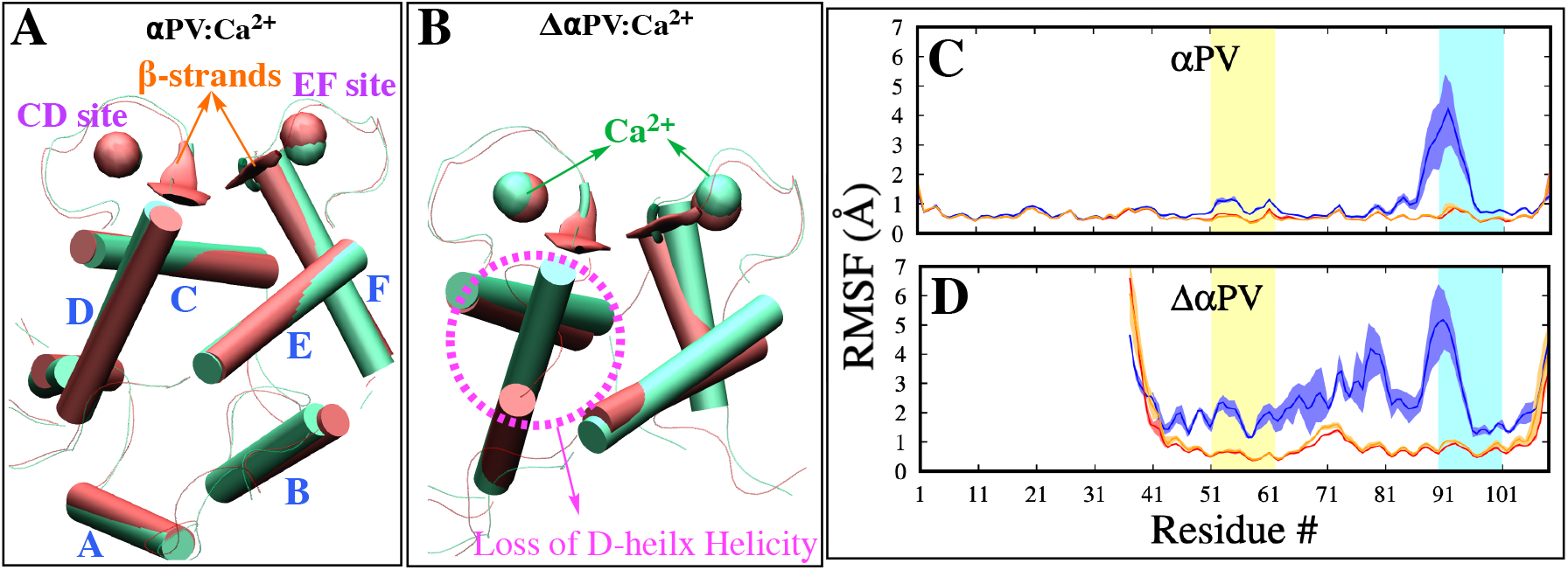
Comparing simulated and experimental structures. **A-B** Superposition of X-ray (green) and MD representative snapshots (red) of WT (A) and Δ*α*PV (B) Ca^2+^ bound systems. **C-D** RMSF vs. Residue number of WT (C) and Δ*α*PV (D) systems. Apo, Ca^2+^, and Mg^2+^ bound systems are colored blue, red and orange, respectively. Average of three MD trials of each system is shown; standard deviation is shaded. Yellow and cyan regions are CD and EF site loop regions. Residues spanning different parts of the protein are mapped as follows: Helix A (S1–A17), helix B (D25–G34), helix C (A40–I49), CD site loop (L50–E60), helix D (D61–F70), helix E (S78–D90), EF loop (G89–G98), and helix F (V99–E108).

We next evaluated in Fig. 1C–D the per-residue conformational dynamics of the proteins via root mean squared fluctuations (RMSF) to identify potential differences that could correlate with the ion affinity. The RMSFs indicate that bound Ca^2+^ in the holo state expectedly suppress fluctuations relative to the apo state. Similar to observations in other closely related CBPs including troponin C (TnC) [33, 37, 34] and *β*-parvalbumin (*β*PV) [38], the reduction in RMSF upon ion binding is largely restricted to the EF hand loop regions (residues L50–E60 (CD site) and G89–G95 (EF site)) (Fig. 1C), although in Δ*α*PV this impact is seen in the entire protein (Fig. 1D), indicating the role of AB domain in *α*PVs stability. The most significant RMSF changes occur in the EF site loop, as the average maximal peak height in the EF site loop in the apo forms is ≈5 Å compared to ≈1.0Å for the holo. Further, we noted a strong correlation between the apo-state RMSF values and Ca^2+^ affinity in the CD and EF sites, as variants with weaker Ca^2+^ yielded larger root mean squared fluctuations (RMSF) values (i.e., the Δ*α*PV) relative to those with stronger affinities (i.e., the WT *α*PV). We did not observe any statistical difference between the Ca^2+^ and Mg^2+^ bound forms (Fig. 1C,D). Overall, these data indicate strong correlation between the extent by which RMSFs are reduced upon Ca^2+^ binding and experimentally-measured binding affinities.

### 3.2 Ranking *α*PV variants’ binding affinities

Our simulations suggest that ion binding reduces the RMS fluctuations intrinsic to the *α*PV apo state. Here we used the end-point thermodynamics method, MM/GBSA, to determine the energetic bases for these changes that may contribute to high Ca^2+^ and Mg^2+^ affinity. Although MM/GBSA is generally recognized to be less accurate than free energy methods like thermodynamic integration and free energy perturbation, it can provide semi-quantitative estimates to help interpret structural trends in molecular simulations [39, 40, 41].

The Generalized Born Surface Area approach used for the MM/GBSA scoring was developed as a fast alternative to computing electrostatic energies from Poisson-Boltzmann (PB) theory [42]. However, the standard PB theory suffers from inaccuracies when applied to highly charged systems [43], such as the Ca^2+^-bound EF-hand loops we considered here. Therefore, we determined if omitting the Ca^2+^/protein binding interaction terms would be more accurate, by assessing the ‘reorganization energy (RE)’ of the protein upon binding an ion via the molecular mechanics score alone. We define RE as the difference in molecular mechanics scores (MM scores) of the ‘isolated protein’ in its holo state versus that of its apo form. The estimated REs are shown in the Table 1. To compare the truncated and the WT variants, we report ΔΔ*G* ≡ Δ*G_i_* − Δ*G_j_*, which cancels out contributions from the free Ca^2+^ s’ solvation energy.

**Table 1:** Estimation of protein reorganization energies

| System | $G_{RE}^{Calc}(\text{holo})$ | $\Delta G_{RE}^{Calc}$ | $\Delta\Delta G_{RE}^{Calc}$ | $\Delta\Delta G^{Expt}$ |
| --- | --- | --- | --- | --- |
| $\alpha\text{PV}:\text{Ca}^{2+}$ | $-1672.66 \pm 1.19$<br>$-1932.35 \pm 7.94^*$ | $128.78 \pm 2.28$ | $0.00 \pm 3.24$ | $0.0 \pm 0.08$ [18, 44, 16] |
| $\Delta\alpha\text{PV}:\text{Ca}^{2+}$ | $-1257.09 \pm 0.34$ | $128.52 \pm 0.48$<br>($415.57 \pm 4.00$)<br>($675.26 \pm 0.59^*$) | $-0.26 \pm 0.76$ | $9.5$ [1] |
| $\alpha\text{PV}:\text{Mg}^{2+}$ | $-1552.78 \pm 0.69$ | $248.66 \pm 2.07$ | $0.0 \pm 2.93$ | $0.0 \pm 0.14$ [18, 44, 16] |
| $\Delta\alpha\text{PV}:\text{Mg}^{2+}$ | $-1134.73 \pm 0.41$ | $250.88 \pm 0.53$<br>($418.05 \pm 0.61$) | $2.22 \pm 0.82$ | nb<sup>a</sup> [1] |

The computed 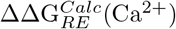 Δ*α*PV compared to the WT is −0.26 ±0.76 kcal/mol. This suggests that the forming of the holo state in Δ*α*PV incurs a smaller thermodynamic penalty than the WT. However, the apo Δ*α*PV structures we simulated remained partially folded and thus do not represent the fully unfolded state as suggested by circular dichroism experiments [1], thus the 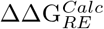 of Δ*α*PV was not meaningful. Therefore, in subsequent analyses of Δ*α*PV we only consider the energy decomposition of its holo state (i.e., 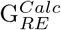 Table 1). The 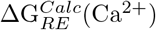 of the Δ*α*PV compared to the WT is 415.57±0.55 kcal/mol, indicating that forming a holo state of Δ*α*PV in the presence of Ca^2+^ incurs a greater reorganization penalty than the WT. This is consistent with the Δ*α*PV exhibiting lesser Ca^2+^ affinity than the WT (≈9.5 kcal/mol), however, we are not indicating that these two estimates are comparable. We observed roughly similar trends for Mg^2+^ as well, i.e., binding of Mg^2+^ to Δ*α*PV incurs greater reorganization penalty relative the WT 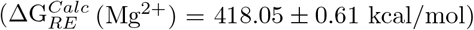. Importantly, we note that 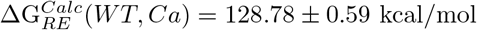 versus 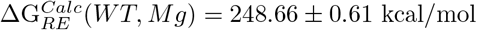. These energies suggest that binding Mg^2+^ manifests an even greater reorganization penalty relative to Ca^2+^ that would disfavor the binding of magnesium. Studies reported that Mg^2+^ does not bind to the Δ*α*PV [1], therefore an experimental ΔΔG^*Expt*^ for the binding of Mg^2+^ is not provided in Table 1.

### 3.3 Ca^2+^ affinity determinants within the EF hands

Since binding occurs within the EF hands, it was expected that changes in binding free energy could be explained by differences in the binding loop conformations that coordinate Ca^2+^ and Mg^2+^. We therefore measured the radial distribution of protein oxygens around bound ions in the CD site to assess the loops’ coordination of Mg^2+^ and Ca^2+^ for the protein (Fig. 2). We observed that the radial distribution maxima for Δ*α*PV occurred at smaller Ca^2+^/oxygen distances (≈2.2 Å) relative to that of the WT (≈2.4 Å) (Fig. 2A). However, the cumulative oxygen density (Fig. 2C) obtained by integrating the radial distributions with respect to distance (Fig. 2A) indicate that within 3 Å, the WT has a near optimal coordination number of eight oxygens, while Δ*α*PV cumulative density reflect seven bound oxygens. We observed similar trends in the EF site, except that the corresponding cumulative distributions indicate WT *α*PV coordinates seven oxygens within 3.0 A of Ca^2+^, whereas for the Δ*α*PV only six are coordinated (Fig. 2B,D). We additionally assessed the number of water oxygens bound to each ion (see Section S8.3). We demonstrate that a coordinated water helps to offset this loss of coordination in the EF site (Fig. 2F,H). The combined contributions of the coordinating amino acid and water oxygens yield similar trends in the total cumulative density of bound oxygens in the EF site for Ca^2+^ (i.e., eight vs. seven in WT*α*PVand Δ*α*PV, respectively (Fig. 2J)) as compared to the CD site (Fig. 2I). Note that no water coordination was observed in the CD site (Fig. 2E, G) within 3 Å of Ca^2+^. As anticipated the weaker Ca^2+^ binding variants (i.e., Δ*α*PV) coordinate fewer oxygens relative to the WT. Overall, these data suggest that the oxygen coordination number does correlate with affinity and that the AB domain is required for the protein to coordinate with maximal efficiency with the binding ions in the EF hands and this demonstrate that the effect of AB domain is allosteric.

**Figure 2:**
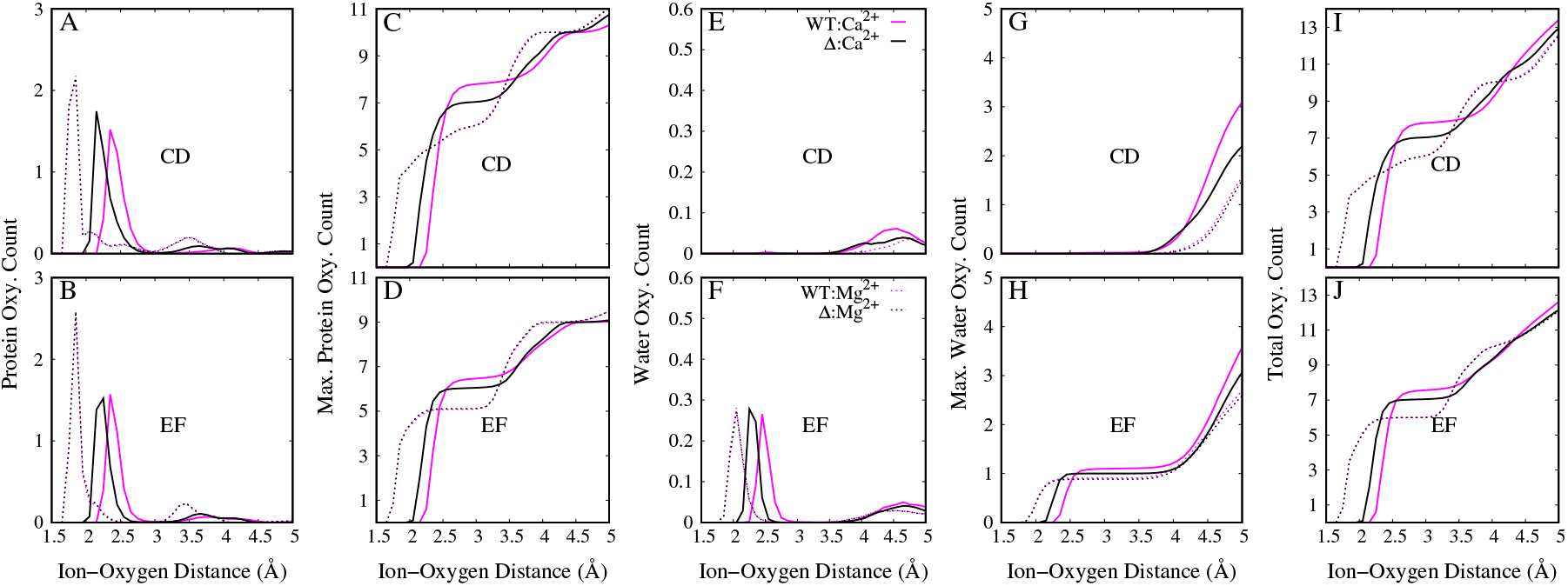
Protein and water co-ordination with the ions in the CD and EF sites. **A, B** Radial distribution of protein oxygens in the CD (A) and EF (B) sites. **C, D** Maximal amino acid oxygen count in CD (C) and EF (D) sites. **E, F** Radial distribution of water oxygens in the CD (E) and EF (F) sites. **G, H** Maximal water oxygen count in CD (G) and EF (H) sites. **I, J** Total oxygen count (i.e., the sum of water and amino acid oxygen count) in CD (I) and EF (J) sites. WT *α*PV and Δ*α*PV are colored magenta and black, respectively.

### 3.4 Ion affinity determinants beyond the EF hands

In the previous sections we have demonstrated that MD-predicted differences in the EF-hand coordination and associated reorganization energies among the variants strongly correlate with experimental measurements. Additionally, we found that point mutations also impart changes in the overall structure of the protein (Fig. 1B-D), as well as its dynamics (Fig. 1F-H). We therefore investigated properties of the entire protein that could further contribute to the range of affinities reported for *α*PV variants, including desolvation, helix bundle ‘packing’, and secondary structure.

Graberek et al. [45] speculated that waters coordinated to solvated Ca^2+^ are released into the bulk medium as the ion binds protein. These liberated waters increase the configurational entropy of the bulk medium and thereby lower the free energy of the system. In this vein, we investigated analogous changes in the solvation of Δ*α*PV that occur during Ca^2+^ binding. In Fig. 3C, we report the number of waters within 4.0 Å of the protein as an indicator of solvation. We observe a reduction (green bars) of ~50 bound waters in the holo states of both WT and Δ*α*PV variants relative to their respective apo states. We anticipate that this number would be far greater in the case of Δ*α*PV if the completely unfolded apo state was considered, which was not captured by our MD simulations. A similar trend was observed in the Mg^2+^ bound systems, and no significant differences were found relative to the Ca^2+^ bound systems. These trends in water loss are similar to those observed for the protein’s radius of gyration (Fig. 3A). The changes in water solvation appear to arise from a reduction in the protein’s solvent accessible surface area (SASA) upon ion binding (Fig. S3). Overall, we observe increased dehydration upon ion binding in both WT and Δ*α*PV variants.

**Figure 3:**
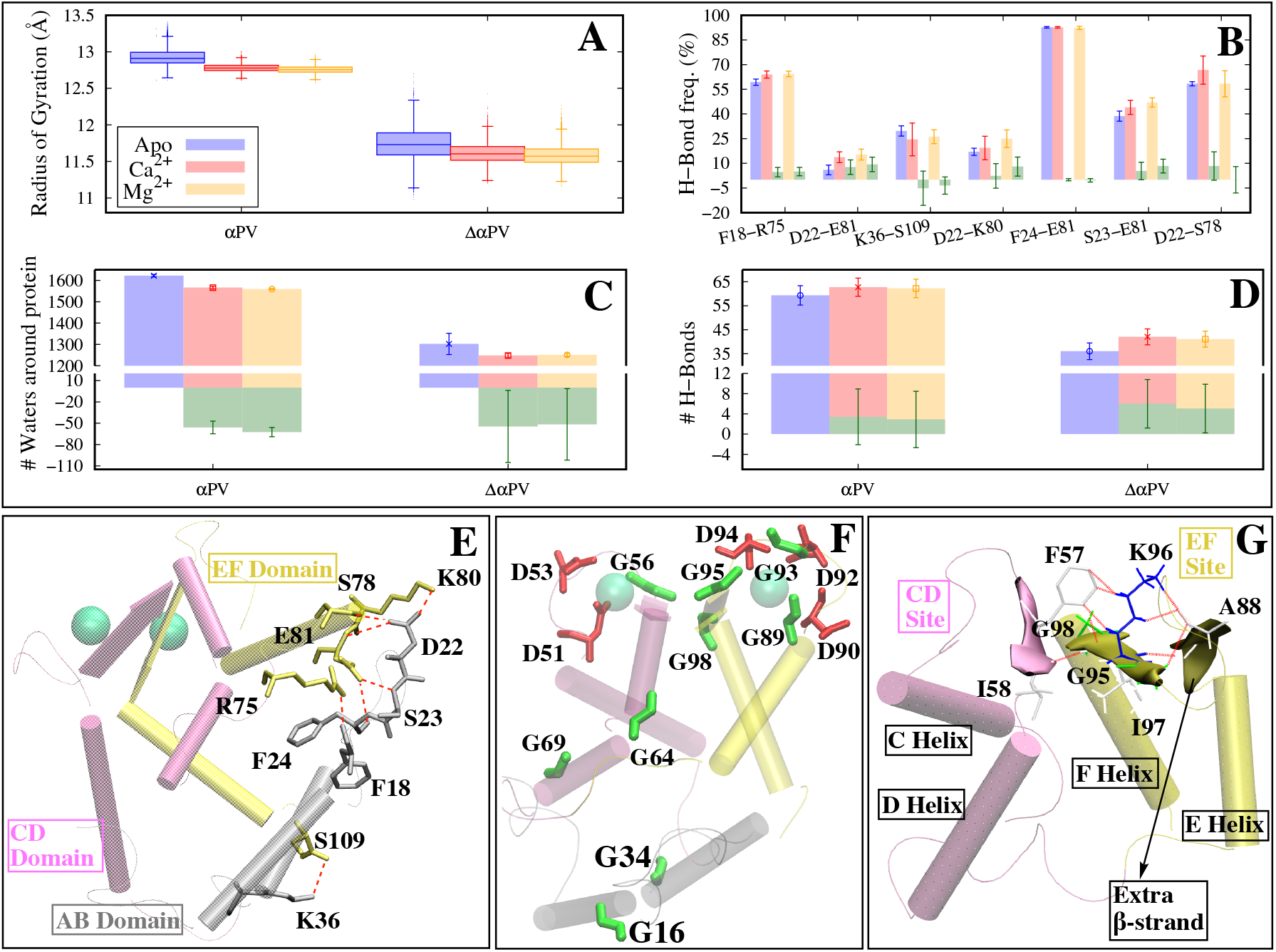
**A** Comparison of radii of gyration (R_*g*_). All three MD trails of a system are plotted as a single box plot. Boxes represent the 25th to 75th quartile (i.e., the interquartile range where 50% of data is concentrated), bottom and top whiskers are minima and maxima, and the horizontal lines in the boxes represent the median, and the points above and below the whiskers are outliers. **B, E** Hydrogen bonds that are formed between the AB domain and remainder of the protein in the holo WT *α*PV. These are absent in the Δ*α*PV since there is no AB domain. Hydrogen bonds are shown as broken red lines (E). **C** Waters within 4 Å of protein. Average and standard deviation of three MD trials are shown. **D** Total number of protein-protein hydrogen bonds. Mean and standard deviation estimated from three MD trials of each variant is shown. Green bars in panels C and D depict the difference of holo and apo states of each respective variant (ΔW (C) and ΔHB (D)). **F** Glycines (green) and aspartic acids (red) in WT *α*PV. **G** Extra *β*-strand formed in EF site loop (colored yellow) in the apo Δ*α*PV. Extra *β-β* hydrogen bonds that are formed in EF site loop region are shown as broken red lines and the corresponding residues are shown as sticks. AB, CD and EF domains are colored grey, magenta and yellow, respectively, in panels E-G.

In Fig. 3D we note that the total number of sidechain-sidechain hydrogen bonds in Δ*α*PV is significantly smaller (≈35) compared to the intact systems (≈60) as would be expected from the loss of its AB helices. Interestingly, the change in hydrogen bonds (ΔHB), i.e., the difference in hydrogen bonds between holo and apo systems, for Δ*α*PV is greater than zero (≈6), which suggests Ca^2+^ plays an important role in increasing the connectivity of the hydrogen bond network within the CDEF helices. We expect that the actual ΔHB would be far greater if we had assumed an completely unfolded conformation for the apo state. From this set, we identified seven hydrogen bonds spanning the AB/CDEF interface that underwent changes in bonding frequency during ion binding (see Fig. 3B). Bonds that exhibited greater contact frequency upon Ca^2+^ binding include F18/R75, D22/E81, D22/K80, D22/S78, S23/E81, and F24/E81, while K36/S109 had fewer contacts. Although nearly all of these reflected increases in bonding frequency in the holo relative to apo states, those changes were generally minor. In other words, the hydrogen bonding network was modestly ‘tightened’ relative to its apo configuration, as opposed to undergoing dramatic changes in the hydrogen bond network. These structural changes culminated in a net gain of ≈18 kcal/mol upon deleting the AB domain from *α*PV (Table 1 and Fig. S4), as assessed using reorganization energies for residues K37/M37 to E108 in *α*PV. Hence, it is apparent that these hydrogen bonds between AB/CDEF helices in the WT *α*PV are favorable and undoubtedly help to stabilize the folded state.

Given the general increase in hydrogen bonding upon metal binding as discussed in the last section resulted in changing of *α*-helix, *β*-sheets and their corresponding energies. *α*-helices represent the most abundant secondary structure in *α*PV (≈45–55%) (Fig. 4A). Our MD simulations indicate that binding ions increases the *α*-helical content in both the WT and Δ*α*PV (≈3–6%), with Ca^2+^ to a greater extent than Mg^2+^ (≈1–4%) (Fig. 4A). We also demonstrate that truncating the AB domain reduces the overall helical content, particularly within the D-helix. Specifically, the greatest impact on the helicity of the D-helix was restricted to the N-terminal end (spanned by residues E60-S65), ~55% in Δ*α*PV vs. “-90% for the WT (Fig. 4C). The corresponding reorganization energy from MM scores indicated that the D-helix was more thermodynamically favored in the WT holo *α*PV relative to Δ*α*PV (≈−149 vs. ≈−137 kcal/mol for Ca^2+^ bound *α*PV and Δ*α*PV, respectively (Table S4)). For the E helix, differences were not apparent (Table S5). These data indicate that the AB domains maintain a high degree of *α*-helicity in the D-helix upon ion binding, which likely contributes to high Ca^2+^ binding affinity.

**Figure 4:**
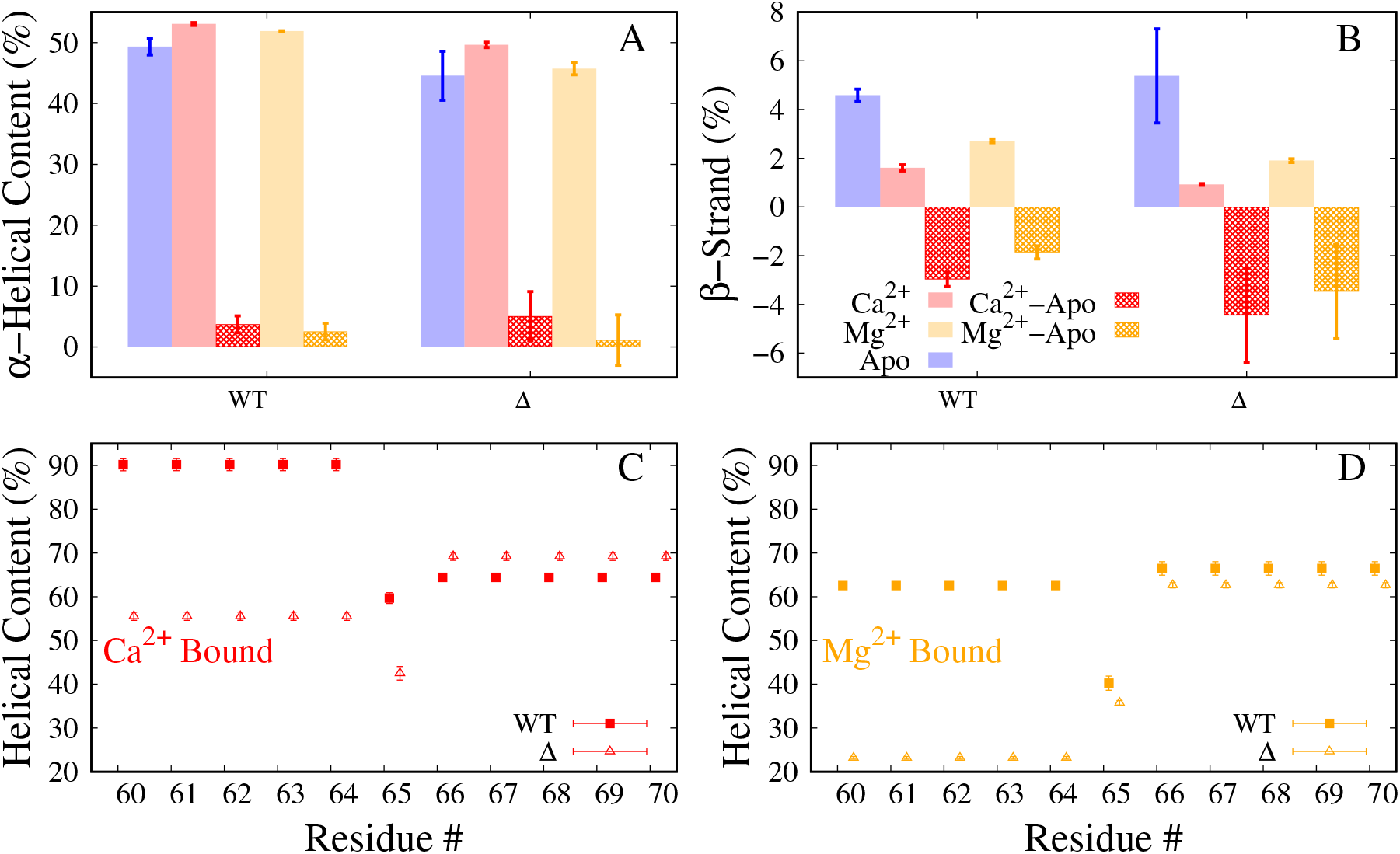
Secondary structural elements compared. **A-B** *α*-helicity (A) and *β*-strand (or extended configuration) (B) of the whole protein. Average and standard deviation of three MD trials of each variant (taken from the means of each MD trial) is shown. The difference between the holo and apo states is also plotted as grid bars. Only C_*α*_ atoms are considered for this calculation. **C-D** *α*-helicity of the D-helix residues compared between the Ca^2+^ (C) and Mg^2+^ (D) bound systems. Average and standard deviation of three MD trials is shown.

## 4 Discussion

The goal of this study was to investigate the role of AB domain in modulating the affinity of Ca^2+^ and Mg^2+^ for *α*PV. To accomplish these goals, MD simulations and thermodynamic analyses of ion binding were conducted of WT *α*PV and Δ*α*PV, truncated variant. We discuss the factors that contribute to the ion binding properties within Δ*α*PV by relative significance

### 4.1 Rank-ordering *α*PV variants according to binding affinity

One of the goals of this study was to elucidate the thermodynamic factors governing the variations in affinity among variants of *α*PV. Explicit free energy techniques including thermodynamic integration, umbrella sampling and adaptive biasing force [46] have been used for rigorous estimates of Ca^2+^ affinity for a variety of Ca^2+^-binding proteins [47, 23], however, it is difficult to decompose free energies predicted by these approaches into per-residue contributions. This limits our ability to implicate specific structural and dynamical differences between variants to their respective ion affinities. Therefore, we have employed an endpoint method, MM scores [46] to rank-order variants by their relative affinity differences. However, including Ca^2+^/protein interactions yielded energies inconsistent with the *α*PV proteins’ experimentally measured Ca^2+^ affinities, which we attribute to shortcomings in the evaluation of polar solvation energies. Therefore, we rather show that reorganization energy of forming the holo state upon ion binding relative to the apo rank-ordered the variants correctly. We have successfully demonstrated the usefulness of this strategy in a previous study [30]. Since the apo Δ*α*PV structures we simulated remained partially folded and thus do not represent the fully unfolded state as suggested by circular dichroism experiments [1], therefore using 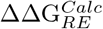 to compare and to rank the variants was not meaningful. Rather, we only considered the energy decomposition of the holo states of the variants (i.e., 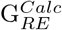 in Table 1). The 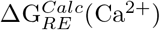 of the Δ*α*PV compared to the WT was 415.57 ±0.55 kcal/mol, indicating that forming a holo state of Δ*α*PV in the presence of Ca^2+^ incurs a greater reorganization penalty than the WT. This is consistent with the Δ*α*PV exhibiting lesser Ca^2+^ affinity than the WT (≈9.5 kcal/mol). We observed roughly similar trends for Mg^2+^ as well, i.e., binding of Mg^2+^ to Δ*α*PV incurs greater reorganization penalty relative the WT 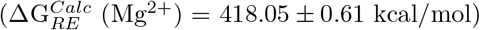. Importantly, we note that 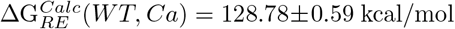 versus 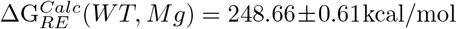. These energies suggest that binding Mg^2+^ manifests an even greater reorganization penalty relative to Ca^2+^ that would disfavor the binding of Mg^2+^.

### 4.2 Contributions of EF-hand loop and co-ordination shell to altered Ca^2+^ binding

The CD and EF domains are the primary binding sites for Ca^2+^ binding in *α*PV proteins and form the helix-disordered loop-helix EF-hand motif. The disordered loop reorganizes to coordinate its amino acid oxygens with Ca^2+^, which manifests in the reduced RMS fluctuations in both WT and Δ*α*PV upon ion binding predicted by our MD simulations. The most significant reductions were apparent within the EF site loop (Fig. 1F-I), and we anticipate this is partly due to the increased presence of glycines and aspartic acids in EF hand loop. There are four glycines and three aspartic acids in the EF hand loop, where as in the CD hand loop there are one and two, respectively. Glycine is categorized as a flexible residue among other aminoacids and is also known as a helix breaker; on the other hand, aspartic acid is relatively considered a flexible residue, particularly compared to glutamic acid [48, 49]. We propose that the knocking of these glycines from the EF site loop transforms it into a CD like site in *α*PV, atleast to some extent. Interestingly, we observed that the amplitude of the apo state RMSFs in the Ca^2+^ binding sites anti-correlated with the variants’ ion affinities, with the largest RMSFs reported for Δ*α*PV and smallest for Δ*α*PV(Fig. 1F-I and Fig. S2). Especially for the CD-loop sites, The calculated MM scores 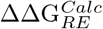 energies for the both the CD and EF sites clearly indicated that the conformational changes upon Ca^2+^ binding yielded more favorable values for WT relative to Δ*α*PV (Table S2). Overall, we show that a decreasing apo state stability in intact PVs is associated with decreased Ca^2+^ affinity.

It is obvious to interpret differences in affinities among the variants based on the properties of Ca^2+^ binding loop, as studies have argued that the position and charge of amino acid substitutions can dramatically enhance Ca^2+^ affinity [50]. For instance, it is expected that increasing the number of oxygens bound to Ca^2+^ could improve the affinity, as has been rationalized for selective binding of Ca^2+^ over Mg^2+^ based on bidentate glutamate binding relative to monodentate [45, 51]. Our data indicate that the number of protein oxygens coordinating the ion is correlated with their affinity, i.e., the number of coordinating oxygens is greater in the WT (a strong binder) compared to the Δ*α*PV (a weak binder) (WT:Ca^2+^ vs. Δ*α*PV:Ca^2+^ is 16 and 12, respectively (Fig. 2 I,J). Our analyses firmly indicate that loop/ion interactions represent significant contributions to the MM scores-estimated ΔΔG s relative to dthe Δ*α*PV (Table S2). We used MM scores energies to quantify the contributions of oxygen and water coordination with Ca^2+^ (Fig. S4 and Table S2) and we found that Δ*α*PV is thermodynamically unfavorable (by ≈24 kcal/mol when both CD and EF sites were combined), compared to the WT, though these calculations neglected stabilizing Ca^2+^/protein interactions. This provides evidence that coordination shell is a major factor that contributes to increased Ca^2+^ affinity in *α*PV and that the AB domain is required for the optimal ion and oxygen coordination.

### 4.3 Structural contributions beyond EF-hand loop in determining Ca^2+^ binding

The coupling between EF-hand binding and allosteric changes in *α*PVs helical bundle organization is exemplified in Ca^2+^ binding proteins such as calmodulin [52, 53], TnC [54, 55] and S100A1 [56, 57, 58]. We show that the enhanced binding of Ca^2+^ to *α*PV protein correlates with several structural changes beyond the EF-hand loop region. The predominant changes caused by ion binding include: (1) reduced hydration shell around the protein, (2) increased hydrogen bonding, (3) increased *α* helicity, particularly in the D and E-helix regions, and (4) reduced *β*-strand content between the CD and EF loops. We discuss each of these factors in order of importance.

We found that the subtle conformational changes induced by ion binding manifested in reduced hydration of holo *α*PV compared to apo (≈20) and to a similar extent in Δ*α*PV (Fig. 3C). This desolvation has been suggested to be a significant factor in driving ion binding in Ca^2+^-selective proteins [51, 45], which can be rationalized by the liberated solvent increasing the configurational entropy of the system. The displacement of loosely bound waters in the CD and EF loops of the apo states upon binding Ca^2+^ represents a substantial portion of the total number of waters liberated in the holo state. It is unclear how much each liberated water contributed to the Ca^2+^ affinity, although one report suggested that a Glu to Asp mutation reduced Ca^2+^ affinity by five-fold estimated based on 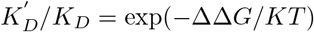 due to an extra coordinated water [59].

The dampened RMSF values and dehydration of holo-state proteins relative to apo states reported in our simulations implicate changes to the organization of the proteins’ helix bundles. To highlight these effects, we measured the net number of hydrogen bonds formed during the apo to holo state transition, as these bonds would be expected to yield larger free energy contributions than the corresponding interactions between apolar residues. We found an increase in the number of intraprotein hydrogen bonds in the holo states of both WT *α*PV and Δ*α*PV variants (≈4–7; Fig. 3D), which were correlated with the dehydration discussed in the previous section. Increasing hydrogen bonds was observed in the holo versus apo states, with similar changes noted for Ca^2+^ and Mg^2+^.

Ca^2+^ binding proteins commonly exert regulatory control through interfacing with target proteins. The binding of the target peptide is often associated with an increase in the apparent Ca^2+^ affinity of the CBPs and increased thermostability [1, 60]. The CD and EF helix-loop-helix motifs in the parvalbumin class of CBPs bear high structural similarity to the globular domains of calmodulin (CaM) and TnC. Uniquely, the AB domains in PV assume a position similar to that of the target-bound protein-protein interface in CaM and TnC, where it binds its CD and EF domains through seven hydrogen bonds. In this regard, the AB domains in PV serve as an ‘endogenous’ peptide that confers thermostabilty and enhanced Ca^2+^ affinity. Indeed, experimental studies support that this AB domain is essential for stabilizing the apo-state *α*PV into its folded form and protects the *α*PV hydrophobic core from hydration, whereby Ca^2+^ affinity is enhanced [1]. Interestingly, a truncated PV variant that lacks the AB domain maintains the ability to bind Ca^2+^ albeit with significantly reduced affinity, whereas a similar truncation in the structurally-similar calbindin 9K is incapable of binding the ion [60]. Our simulation results confirm that the intact *α*PV has increased hydrogen bonding and reduced bundle mobility as evidenced by RMSF calculations, helicity, and dehydration upon Ca^2+^ binding relative to the truncated variant. These differences highlight the importance of the AB domains in defining *α*PV’s Ca^2+^ binding properties. Supporting these observations, in the holo state there is a loss of favorable reorganization energy of the CDEF domain in the truncated variant compared to the intact one (Table 1). Despite this loss of stability, the Δ*α*PV holo state remains folded in the presence of Ca^2+^ as supported by the Thepaut *et al* experiments [1]. Nonetheless, even though the Δ*α*PV holo structure is folded, it presents high RMSF values that suggest instability within the helix bundle relative to the intact *α*PVvariants, in addition to reduced D-helicity and enhanced *β*-sheet character that may impair Ca^2+^ binding. While we were unable to predict the unfolded apo state indicated by Thepaut *et al*, our simulations suggest that the Δ*α*PV apo structures were unfolding and highly labile relative to the intact *α*PV structure. From this perspective, the presence of the AB domains may amortize the thermodynamic cost of folding the protein upon binding Ca^2+^, which would otherwise reduce the *α*PV apparent Ca^2+^ affinity. In support of this interpretation, lysine-to-asparagine mutations of the AB domain sites A8 and H26 in PV constructs modeled after Antarctic fish were found to increase Ca^2+^ binding affinity and were attributed to forming additional hydrogen bonds within the AB or between AB and CDEF domains [61]. Overall, our observations highlight the importance of AB/CDEF domain interactions for maintaining high Ca^2+^ binding affinity.

As with many Ca^2+^ binding proteins, *α*PV is predominantly composed of *α*-helices that undergo displacements and changes in folding upon binding by Ca^2+^ [62, 63, 64, 51]. We observed a consistent trend in our simulations of increased *α*-helicity in the ion-bound holo states relative to apo (≈5%; Fig. 4A), with greater gains demonstrated for Ca^2+^ relative to Mg^2+^, although not significant. This observation is consistent with observations from Laberge *et al*., who through ultraviolet circular dichroism spectra and picosecond MD simulations of cod PV [65] showed that removal of Ca^2+^ reduced *α*-helical content. Our simulations indicate that the majority of the gain in *α*-helical character stems from refolding of the N-terminal end of the D helix. At these termini, glutamic acids at the 59th and 62nd sequence positions are reoriented by Ca^2+^ as the ion binds the carboxylic acid side chains. This process increases the helicity of residues E59 through G64 to 90% (Fig. 4 C, D). In contrast, we observed an increase in helicity in the C-terminal end of the E helix, particularly for the residues T84-D90 (Fig. S6). Interestingly, our Δ*α*PV simulations reflected a very low (55%) degree of helicity formed in the D-helix N-terminus, which suggests the AB domains likely play an important role in stabilizing the Ca^2+^ binding loop, which in turn may increase its affinity for Ca^2+^. In addition, the D-helix N-terminus has a lesser degree of *α*-helicity in the Mg^2+^-bound state (65%). It has been speculated that this difference may be the consequence of the 12th position glutamic acid (E62) promoting bidentate oxygen coordination for Ca^2+^ relative to monodentate for Mg^2+^ [51, 66]. No such differences were noted in the E-helix between the Ca^2+^ and Mg^2+^ ions (Fig. S6). Altogether, our data indicate that increasing the *α*-helicity particularly in the N-terminal region of helices D and E, may play an integral role in high affinity Ca^2+^ binding and selectivity against Mg^2+^. These MD observations were supported by reorganization energies of the D-helix residues (Table S4), i.e., the D-helix in Δ*α*PV imposes a ≈11 kcal/mol greater thermodynamic penalty than the WT for binding Ca^2+^ (≈6 kcal/mol in the case of Mg^2+^).

A hallmark of Ca-binding proteins (CBPs) is the pairing of two Ca^2+^-binding EF hand domains via a *β*-sheet, which has been coined as the “EF-hand *β*-scaffold (EFBS)”[51]. It has been reported that *β*-sheets contribute to the Ca^2+^ binding mechanism via coupling the two binding sites through hydrogen bonds and dipole-dipole interactions and that improved Ca^2+^ affinity to the CBPs through positive cooperativity [51, 67]. Our studies support the notion that *β*-sheet character is reduced upon Ca^2+^ binding and to a lesser extent by Mg^2+^. The low Ca^2+^-affinity Δ*α*PV presents a lower degree of *β*-sheet character when bound to Ca^2+^ as well as in the apo state. It is evident from published structural data and simulations that the reduction in *β*-strand content occurs as the residues contributed by each site (F57-E59 and K96-G98) are drawn away from the *β*-sheet interface by their respective coordinated ions. The largest displacements arise from E59 and K96 as they are separated from one another by S55 and D94, respectively, through the formation of hydrogen bonds (Fig. S8). Overall, our simulation data suggest that enhanced *β*-sheet character in the apo state appears to offset the free energy gain upon Ca^2+^ binding. *β* character is correlated with ion affinity, i.e., greater the *β* component greater is the ion affinity (Fig. S7). These observations are supported by MM/GBSA reorganization energies (Table S3).

### 4.4 Determinants of Ca^2+^ versus Mg^2+^ binding

Ca^2+^ binds with greater affinity (≈10 kcal/mol) than Mg^2+^ to WT *α*PV. This in large part can be rationalized by the considerably higher desolvation cost for Mg^2+^ relative to Ca^2+^ and to a lesser extent by the reduced number of protein oxygens bound by Mg^2+^ [51]. Regardless of the origins of the affinity differences, our simulations reveal distinct differences between the holo states of Mg^2+^-bound *α*PV structures relative to those bound with Ca^2+^. These differences include reduced *α*-helicity (Fig. 4A), especially within the D-helix N-terminal region (Fig. 4 C,D), and less significant disruption of the *β*-sheets bridging the EF-hand loops (Fig. 4B). These secondary structure differences are mirrored by the reorganization energies from these corresponding regions in the *α*PV proteins (Table S4 and Table S3). The presence of the AB domains in *α*PV most likely masks such structural differences that would arise for Ca^2+^-bound states relative to Mg^2+^ bound. The AB domains appears to lock *α*PV into an open-like state even in the absence of Ca^2+^, which thereby facilitates the binding of other competing cations such as Mg^2+^. This interpretation is supported by studies from Thepaut *et al* that indicated Mg^2+^ was unable to fold the Δ*α*PV variant that lacked the AB helices. For this reason, the prominence of parvalbumin proteins serving as delayed Ca^2+^ buffers in skeletal and cardiac muscle [68] likely stems from its AB domain facilitating Mg^2+^ binding and thereby competing with Ca^2+^ binding.

## 5 Conclusions

In this study we investigated the manner in which the AB domain of *α*PV regulates the binding of Ca^2+^ and Mg^2+^ to the EF hands, which are distal to the AB domain, as well as its role in selectivity of *α*PV for Ca^2+^ over Mg^2+^. Our approach comprises of simulating WT *α*PV and Δ*α*PV in apo as well as Ca^2+^/Mg^2+^-bound (holo) states via all-atom MD, followed by application of MM/GBSA method. We demonstrate that the MM/GBSA protein ‘reorganization energies’ qualitatively rank ordering both the *α*PV variants (WT and Δ*α*PV) by their experimentally determined Ca^2+^ and Mg^2+^ affinities. We further used reorganization energies from these analyses to implicate structural changes that rationalize the variants’ ion binding affinities. It has been identified that AB domain modulate the preferential binding of Ca^2+^ over Mg^2+^ in an allosteric manner. The major impact of AB domain is particularly seen on the D-helix; absence of AB facilitates the unwinding of the D-helix N-terminus residues 60–65 (~30–35%) in the Ca^2+^ systems, which coordinates the binding of ion in the CD loop. Note that C-terminus of the D-helix interacts with the AB domain. Contrary to our earlier study [30], we find that the number of coordinating oxygens in the EF hand binding sites’ are sufficient to completely rationalize observed changes in affinity.

## 6 Materials and methods

### 6.1 Protein systems

Ca^2+^ bound rat X-crystal structures of *α*PV (PDB: 1RWY [36]) and Δ*α*PV (PDB: 1G33 [1]) were used for the MD simulations. Apo simulations were set up by removing ions from the holo crystal structures. The crystal structures of Mg^2+^ bound *α*PV protein are not yet available, therefore, Ca^2+^ was replaced with Mg^2+^ for the Mg^2+^ bound simulations. System preparation was performed with *tleap* from Amber 16 [69].

### 6.2 Classical MD

All systems were solvated in TIP3P [70] water box with 20Å margin and neutralized with 0.15 M KCl. Each system has about 14,000 waters, 82 neutralizing ions, for a total of ≈44,000 atoms. The systems were parameterized with the amber ff12SB force field [71] with Ca^2+^ and Mg^2+^ parameters were adapted from Li-Merz [25]. The system was subjected to 6000 steps minimization with protein atoms being fixed during the first 1000 steps. The system was then heated under NVT ensemble to 300 K over 0.1 ns using the weak-coupling algorithm [72]. Constraints were introduced on protein atoms with a force constant of 500 *kcal/mol. Å*^2^. The heated system was equilibrated for 1 ns under the NPT ensemble without constraints on the protein. After equilibrium, each system was simulated in triplicate for ≈1 *μ*s under the NPT ensemble at 300 K temperature. A 2 fs time step was used with the non-bonded cutoff set as 10 Å A and the electrostatic interactions were treated with Particle Mesh Ewald (PME) method [73]. The SHAKE algorithm was used [74]. The simulation data of the production run were saved for every 20 ps. The minimization and equilibration were performed using the *PMEMD* module while the production runs were performed using the GPU-accelerated *PEMD.CUDA* module of AMBER16 [69].

### 6.3 Analyses

Two snapshots for each ns of trajectory were considered for the analysis (≈2000 data points for each simulation trajectory (≈1*μ*s) and a total of 36000 data points for all systems combined (6 systems × 3 trials × 2000 data points/ 1*μ*s) ≈36000 data points). Almost all analyses were conducted with VMD [75] and its associated plugins [75, 76], except for the radial distribution functions. For ion bound systems, the radial distribution of chelating oxygens about the cation (either Ca^2+^ or Mg^2+^) was obtained by the *radial* command of *CPPTRAJ* [77]. For calculating hydrogen bonds, the cut-off distance and angle used were 3.5 Å and 30°respectively. RMSDs and RMSFs were calculated by superposing each trajectory frame with the respective x-ray crystal structure. The MMPBSA.py script of AMBER16 was used for the MM/GBSA calculations. The ionic strength for the MM/GBSA calculation was set as 0.15 M with the generalized Born model option set as igb=5. The entropy was neglected during the MM/GBSA calculations. The entire MD trajectories were used for the MM/GBSA calculations. The total reorganization energy of each system was calculated as the sum of all five individual components of the MM/GBSA method, including electrostatic, van der Waals, internal, polar, and non-polar solvation, respectively.

## 7 Supporting Information

Figures S1–S8, Tables S1–S7, and limitations of our strategy are provided in the supplementary information.

## S8 Supporting Information

### S8.1 Figures

**Figure S1:**
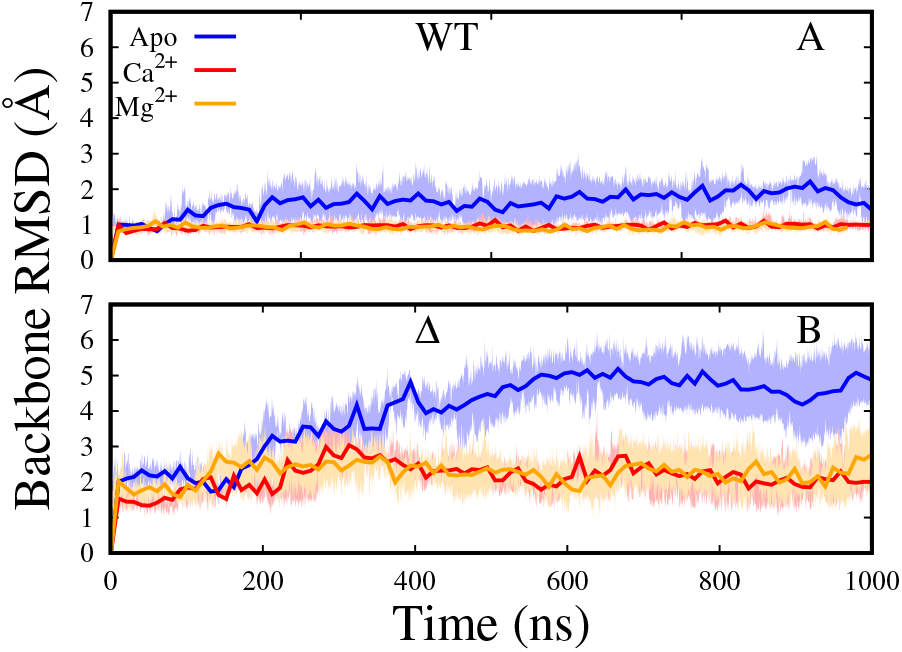
Protein backbone RMSD vs. Time. Apo, Ca^2+^, and Mg^2+^ bound systems are colored blue, red and orange respectively. Dark lines represent the average and shaded regions represent the standard deviation of three MD runs.

**Figure S2:**
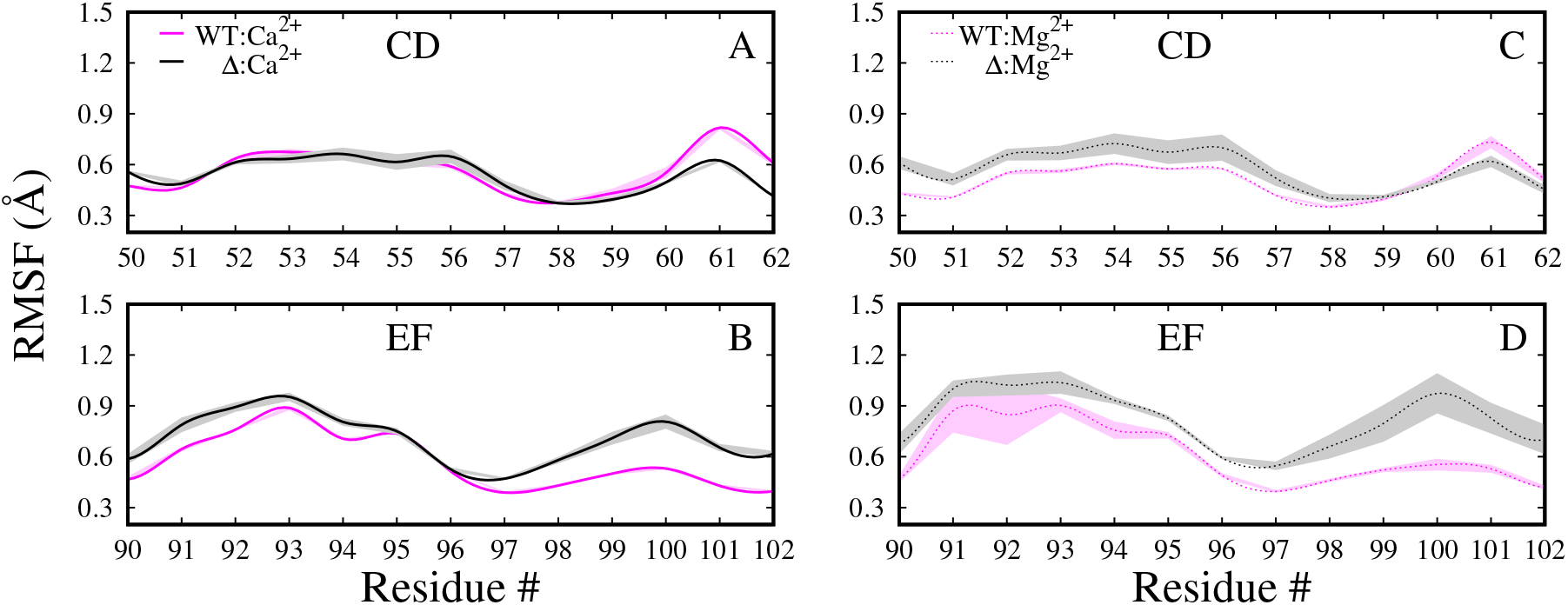
Root mean square fluctuation (RMSF) of the CD (A, C) and EF (B, D) sites of holo systems. Ca^2+^ bound systems are shown in panels A,B and Mg^2+^ bound in panels C,D. Mean and standard deviation of three MD trials are shown.

**Figure S3:**
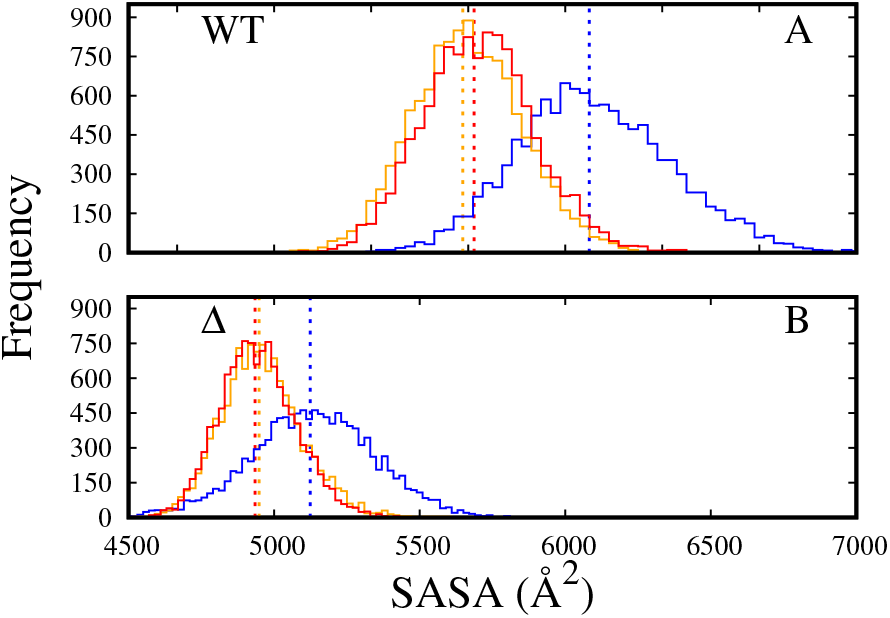
Distribution of protein solvent accessible surface area (SASA). Apo, Ca^2+^, and Mg^2+^ bound systems are colored blue, red and orange respectively. Average of each *α*PV variant are represented by dashed vertical lines.

**Figure S4:**
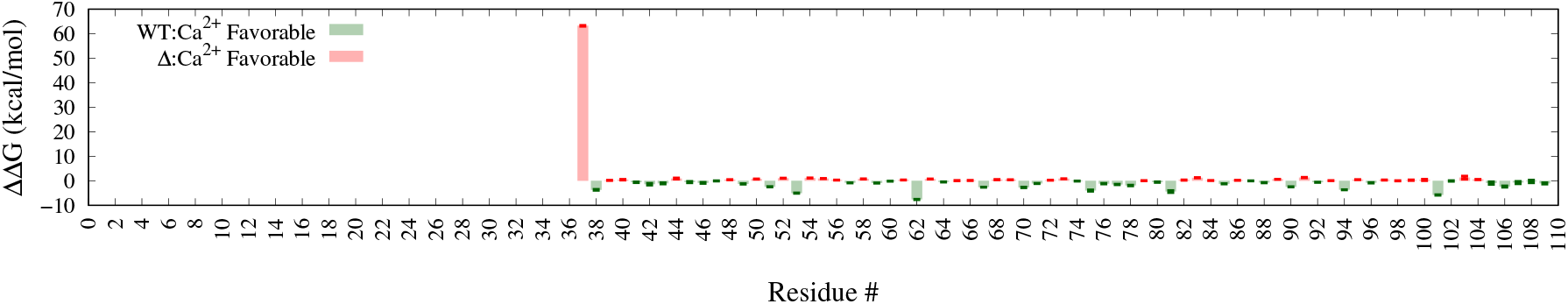
Residue reorganization energies of Ca^2+^ bound systems. The MD predicted Δ*α*PV apo does not unfold as expected from experiment [1], hence, only the free energies of the holo state are compared. See Table S3 and Table S4 for summations of structurally significant regions. Mean and SEM of each residue estimated from three MD trials are shown yes.

**Figure S5:**
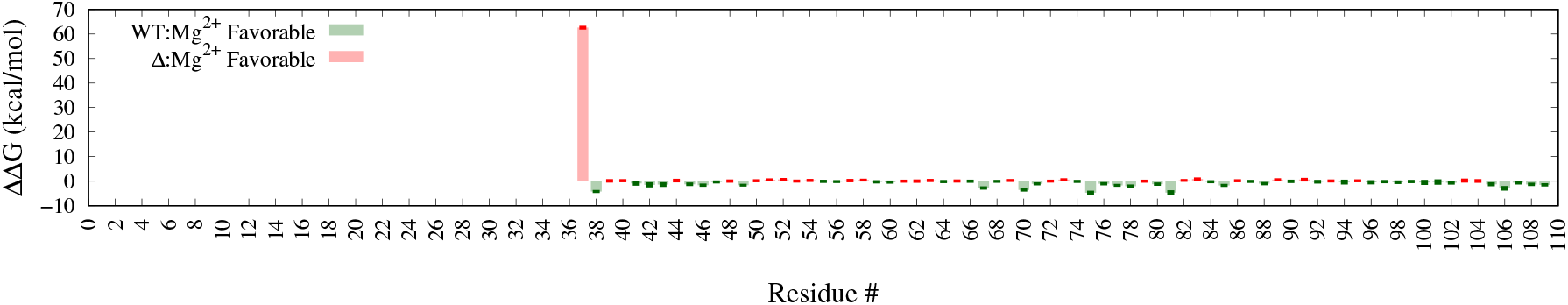
Comparing residue reorganization energies of Mg^2+^ bound systems. Mean and SEM of each residue estimated from three MD trials are shownyes.

**Figure S6:**
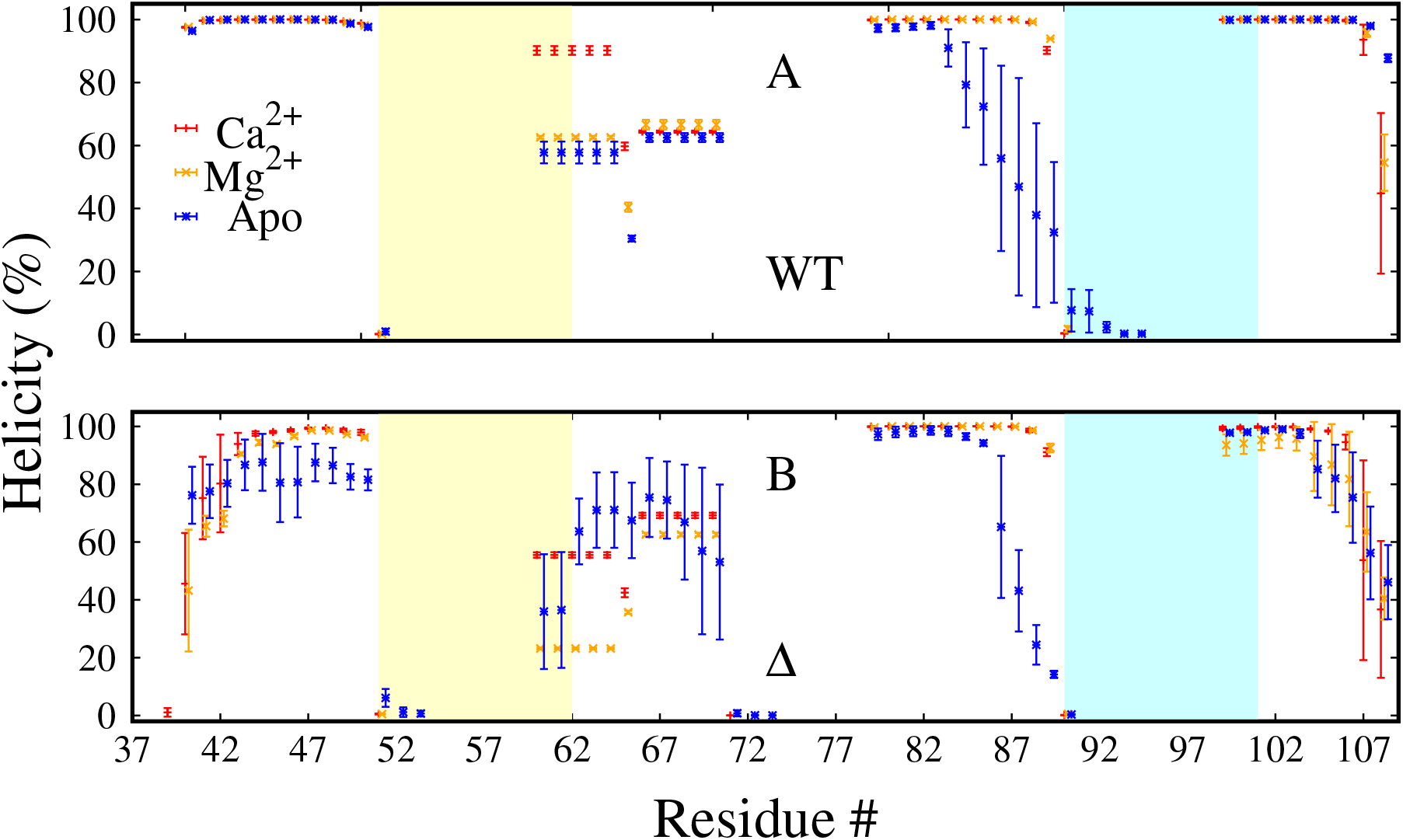
*α*-helicity vs. residue number. Apo, Ca^2+^ and Mg^2+^ bound systems are colored blue, red and orange, respectively. CD and EF site loop domains are colored yellow and blue, respectively. Error bars represent standard deviation estimated from three MD runs.

**Figure S7:**
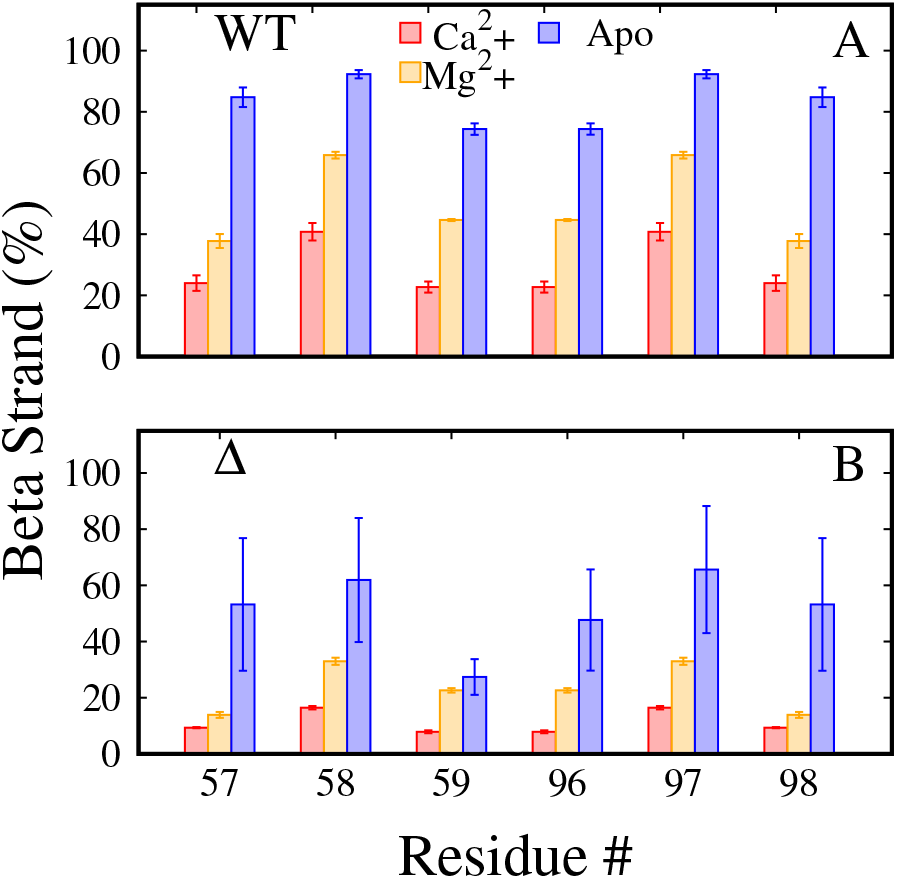
*β*-strand of the six residues (F57–E59 and K96–G98) that form the *β*-β bridge coupling the two ion binding sites. Apo, Ca^2+^ and Mg^2+^ bound systems are colored blue, red and orange, respectively. Error bars represent standard deviation.

**Figure S8:**
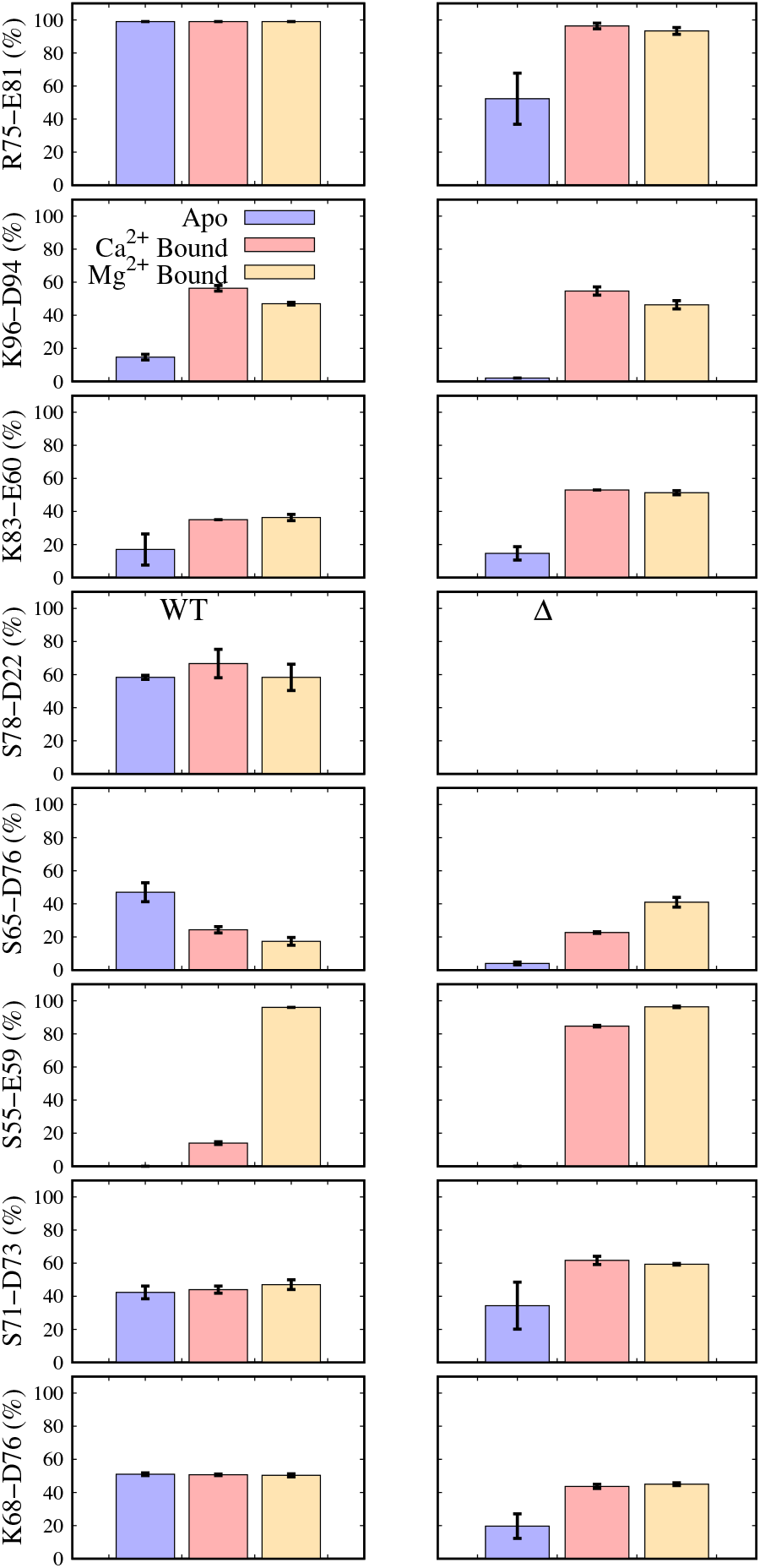
Selective hydrogen bonds (sidechain-sidechain) of interest. Columns 1 and 2 represent *α*PV and Δ*α*PV, respectively. Blue, red and orange represent apo, Ca^2+^ and Mg^2+^ bound systems, respectively. Error bars are standard deviations of three MD trails.

### S8.2 Tables

**Table S1:** Estimated total MM/GBSA energies.

| System | $\Delta G_{Total}^{Calc}$ | $\Delta\Delta G_{Total}^{Calc}$ | $\Delta G^{Expt}$ | $\Delta\Delta G^{Expt}$ |
| --- | --- | --- | --- | --- |
| $\alpha PV:Ca^{2+}$ | $-84.85 \pm 1.27$ | $0.00 \pm 1.79$ | $-22.0 \pm 0.06$ ($-20.5 \pm 0.06$) [18, 19] | $0.0 \pm 0.13$ |
| $\Delta\alpha PV:Ca^{2+}$ | $-154.26 \pm 1.20$ | $-69.41 \pm 3.06$ | $-12.6$ [1] | 9.5 |
| $\alpha PV:Mg^{2+}$ | $-204.61 \pm 1.27$ | $0.00 \pm 1.79$ | $-11.6 \pm 0.04$ ($-10.0 \pm 0.06$) [18, 19] | $0.0 \pm 0.09$ |
| $\Delta\alpha PV:Mg^{2+}$ | $-206.68 \pm 1.20$ | $-2.54 \pm 3.04$ | nb<sup>a</sup> [1] | nb<sup>a</sup> |

$\Delta G_{Total}^{Calc}$ for each system is calculated as the difference of holo and apo state energies. Mean and SEM of all three MD trials of each system are shown. $\Delta\Delta G_{Total}^{Calc}$ is calculated relative energy and $\Delta\Delta G^{Expt}$ is experimental relative free energy, and are calculated as the difference of free energies of the variants with respect to the WT. Experimental values shown in parenthesis are at 5°C and the rest at 25°C. All energies are in kcal/mol. <sup>a</sup>non binder.

**Table S2:** MM/GBSA reorganization energies of CD and EF site residues.

| System | $G_{RE}^{Calc}(CD)$ | $\Delta G_{RE}^{Calc}(CD)$ | $\Delta\Delta G_{RE}^{Calc}(CD)$ | $G_{RE}^{Calc}(EF)$ | $\Delta G_{RE}^{Calc}(EF)$ | $\Delta\Delta G_{RE}^{Calc}(EF)$ | $\Delta\Delta G_{RE}^{Calc}(Total)$ |
| --- | --- | --- | --- | --- | --- | --- | --- |
| $\alpha PV:Ca^{2+}$ | ($-265.86 \pm 0.06$) | $66.96 \pm 0.52$ | $0.00 \pm 0.74$ | ($-248.05 \pm 0.09$) | $43.70 \pm 0.88$ | $0.00 \pm 1.25$ | $0.00 \pm 1.46$ |
| $\Delta\alpha PV:Ca^{2+}$ | ($-254.13 \pm 0.04$) | $79.18 \pm 1.32$<br>($11.73 \pm 0.07$) | $12.22 \pm 1.42$ | ($-237.57 \pm 0.37$) | $55.90 \pm 1.08$<br>($10.48 \pm 0.38$) | $12.20 \pm 1.39$ | $24.42 \pm 1.99$ |
| $\alpha PV:Mg^{2+}$ | ($-190.54 \pm 0.21$) | $142.28 \pm 0.56$ | $0.00 \pm 0.80$ | ($-184.60 \pm 0.82$) | $107.15 \pm 1.20$ | $0.00 \pm 1.70$ | $0.00 \pm 1.88$ |
| $\Delta\alpha PV:Mg^{2+}$ | ($-192.21 \pm 0.04$) | $141.10 \pm 1.32$<br>($-1.67 \pm 0.21$) | $-1.17 \pm 1.44$ | ($-182.21 \pm 1.16$) | $111.26 \pm 1.53$<br>($2.31 \pm 1.42$) | $4.10 \pm 1.95$ | $2.92 \pm 2.43$ |

**Table S3:** MM/GBSA reorganization energies of *β*-sheet residues.

| System | $G_{RE}^{Calc}(\beta)$ | $\Delta G_{RE}^{Calc}(\beta)$ | $\Delta\Delta G_{RE}^{Calc}(\beta)$ |
| --- | --- | --- | --- |
| $\alpha PV:Ca^{2+}$ | ($-30.24 \pm 0.15$) | $12.45 \pm 0.39$ | $0.00 \pm 0.56$ |
| $\Delta\alpha PV:Ca^{2+}$ | ($-28.84 \pm 0.06$) | $14.89 \pm 0.64$<br>($1.40 \pm 0.16$) | $2.43 \pm 0.75$ |
| $\alpha PV:Mg^{2+}$ | ($0.64 \pm 0.17$) | $43.34 \pm 0.40$ | $0.00 \pm 0.57$ |
| $\Delta\alpha PV:Mg^{2+}$ | ($1.22 \pm 0.17$) | $44.96 \pm 0.66$<br>($-1.06 \pm 0.24$) | $1.62 \pm 0.78$ |

Data are included for the $\beta$ -sheet residues (F57–E59, K96–G98). Error bars represent SEM. Holo state energies are shown in parenthesis.

**Table S4:**
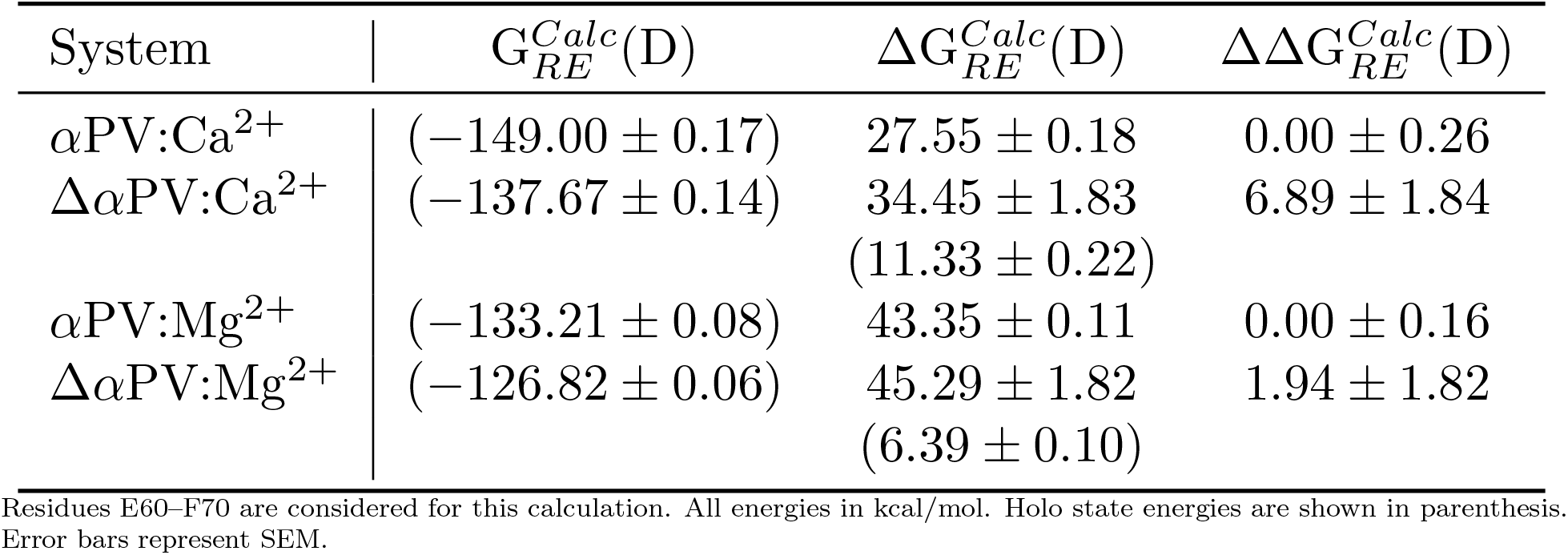
MM/GBSA reorganization energies of the D-helix residues.

| System | $G_{RE}^{Calc}(D)$ | $\Delta G_{RE}^{Calc}(D)$ | $\Delta\Delta G_{RE}^{Calc}(D)$ |
| --- | --- | --- | --- |
| $\alpha PV:Ca^{2+}$ | $(-149.00 \pm 0.17)$ | $27.55 \pm 0.18$ | $0.00 \pm 0.26$ |
| $\Delta\alpha PV:Ca^{2+}$ | $(-137.67 \pm 0.14)$ | $34.45 \pm 1.83$<br>$(11.33 \pm 0.22)$ | $6.89 \pm 1.84$ |
| $\alpha PV:Mg^{2+}$ | $(-133.21 \pm 0.08)$ | $43.35 \pm 0.11$ | $0.00 \pm 0.16$ |
| $\Delta\alpha PV:Mg^{2+}$ | $(-126.82 \pm 0.06)$ | $45.29 \pm 1.82$<br>$(6.39 \pm 0.10)$ | $1.94 \pm 1.82$ |

Residues E60–F70 are considered for this calculation. All energies in kcal/mol. Holo state energies are shown in parenthesis. Error bars represent SEM.

**Table S5:**
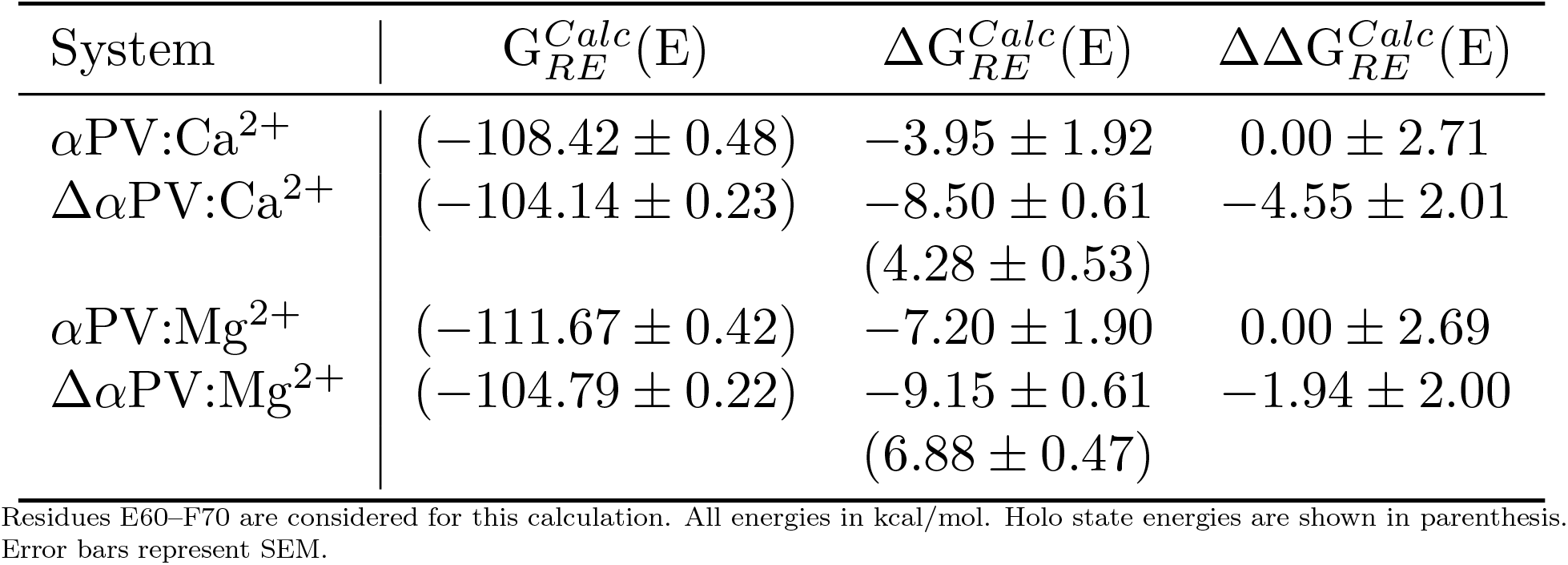
MM/GBSA reorganization energies of the E-helix residues.

### S8.3 Supplementary Results

*Protein backbone RMSDs* Protein backbone RMSDs were calculated by comparing against the respective starting structure. The ion bound forms converged around 1Å (≈2Å for Δ*α*PV) approximately after 200 ns, where as the apo forms converged around 2A after 600 ns (≈5Å for Δ*α*PV). The backbone RMSDs shows that the binding of ions (Ca^2+^ or Mg^2+^) dampens the structural fluctuations of the variants (Fig. S1). While the difference between the apo and ion bound forms is statistically significant in all variants studied, no statistical difference between Ca^2+^ and Mg^2+^ bound forms was observed.

Overall, the structural similarities and converged RMSDs suggest that our simulation protocol can reproduce the conformational ensembles of ion-bound forms and also likely accurately model the apo-forms of the *α*PV variants (except for Δ*α*PV) for which no structural data are available.

*Water coordination of Ca^2+^* We assessed the number of waters coordinated by Ca^2+^ while bound to the CD and EF hand loops (Fig. 2E-H). We relate the cumulative number of protein and water oxygens in panels Fig. 2I-J to determine the total coordination number of Ca^2+^.

*Coordination shell in Mg^2+^ bound systems* In the Mg^2+^ bound systems we demonstrate through radial distribution of protein oxygens that the proteins bind tighter to the Mg^2+^ ion compared to the Ca^2+^. In either binding site, the first oxygen coordination shell is at ≈1.7A for both systems (0.5 A less than the Ca^2+^ bound systems) (Fig. 2A,B). This indicates that protein has to rearrange to interact with the Mg^2+^ (compared to Ca^2+^), which might come at a thermodynamics cost. The number of coordinating oxygens in this CD and EF sites for both systems is five (Fig. 2C,D). Overall, we show that, four fewer protein oxygens are coordinating the Mg^2+^ (in both sites combined) compared to Ca^2+^ bound systems, which partly explains the lower affinity of Mg^2+^ to *α*PV systems compared to Ca^2+^. This is in line with other earlier observations that the Mg^2+^ coordination is lower than Ca^2+^ in PVs [66]. Moreover, the water coordination in the Mg^2+^ bound systems is very similar to the Ca^2+^ bound ones, except that the first hydration shell is closer in the Mg^2+^ bound systems compared to the Ca^2+^ bound systems (≈0.5 A, lower) (Fig. 2E-H).

The score difference between the Δ*α*PV and WT Ca^2+^ bound states was approximately ≈23 kcal/mol (both sites combined), whereas the difference between the Mg^2+^ bound states was ≈3 kcal/mol (Table S2). This data suggests that Ca^2+^ and Mg^2+^ bind less favorable for the truncated species.

*Specific hydrogen bonding interactions* The solvation differences between apo and holo states are driven by conformational changes upon ion binding. We therefore analyzed hydrogen bonds between amino acid side chains to determine if any may help stabilize the holo state. In Fig. S8 we report that for the hydrogen bonds that change between the apo and holo states, a greater number are formed during Ca^2+^ binding as opposed to breaking. This includes the formation of S55-E59 (CD site) and K96-D94 (EF site) hydrogen bonds in the holo forms (Fig. S8). In the Δ*α*PV system, the lack of AB negates the S78-D22 interaction.

In (Fig. 1A-B) we reported structural and dynamic changes between the *α*PVand the variant. Those changes were accompanied by MM scores energies that correlated with the proteins’ Ca^2+^ binding trends and selectivity against Mg^2+^. We therefore investigated properties of the entire protein that could further contribute to the range of affinities reported in *α*PV variant. Firstly, we investigated changes in the solvation of the *α*PV variants that occur during Ca^2+^ binding. This was motivated by speculations in Graberek et al. [45] that waters coordinated to Ca^2+^ are released into the bulk medium as the ion binds proteins. This process can increase the configurational entropy of the bulk medium and thereby lower the free energy of the system. In Fig. 3C, we report the number of waters that are displaced by Mg^2+^ and Ca^2+^ upon binding to the EF-hands. This assessment was based on calculating the integrated radial distribution of waters around the geometric center of EF-hands (Fig. 2). The number of displaced waters is then represented as the difference between the distributions computed for the apo and ion-bound cases. We evaluated this difference at 7.5 Å from the EF hand center of mass, which corresponds to its average distance to Ca^2+^ and a surrounding layer of waters (assuming a 1.4 A radius). We found that, for both Ca^2+^ and Mg^2+^ bindings, the WT displaced more water than the Δ*α*PV. Interestingly, in both cases, Mg^2+^ binding displaced 0.3-0.5 more waters than Ca^2+^ binding.

Across all systems, we observe at least a reduction (green bars) of twenty bound waters in the ion-bound holo states relative to the apo states. Similar trends were observed in Mg^2+^ bound systems, and no significant differences were found relative to the Ca^2+^ bound systems. These trends in water loss are similar to those observed for the proteins’ radius of gyration (Fig. 3A). The changes in water solvation appear to arise from a reduction in the proteins’ solvent accessible surface area (SASA) upon ion binding (Fig. S3). Overall, we observe a general trend of increased dehydration upon ion binding, however these trends does not correlates with the proteins’ affinities for respective ions. It is unclear how much each liberated water contributed to the Ca^2+^ affinity, although one report suggested that a Glu to Asp mutation reduced Ca^2+^ affinity by five-fold due to an extra coordinated water [59], which suggests a ΔΔG of approximately 1 kcal/mol using 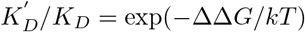.

### S8.4 Limitations

Limitations of our approach include the limited MD simulation length (microsecond), in addition to approximations associated with the MD and MM/GBSA methods (detailed in the review [46]). The time scale of binding, folding and unfolding etc. are usually on the order of milliseconds to seconds [78], compared to our microsecond-length simulations. The other significant limitation is that MM/GBSA is not sufficiently accurate to predict the solvation energies of ions, which is crucial to assess the absolute/relative binding affinities of the ions to *α*PV [46]. We attribute this to the inherent approximations in this end point method, such as the implicit treatment of the solvent, neglect of entropy, and the sensitivity of the method to the solute dielectric constant and the challenges in accurately modeling the non-polar contribution to the solution free energy [40, 79, 80]. Force field deficiencies might also contribute to the inaccuracies in the binding affinity estimation, particularly those pertaining to describing interactions with divalent ions like Ca^2+^ [81, 25].

We believe Δ*α*PV was somewhat problematic because this simulation was started from the completely ordered holo structure by removing Ca^2+^, since the apo state crystal structure of the Δ*α*PV was unavailable; experimental studies report that the Δ*α*PV apo structure is completely disordered [1]. However, in our simulations we only observed partial unfolding of the structure, which warrants more simulation time that is beyond the scope of this study.

## Acronyms

ΔαPV: truncated *α*-parvalbumin. 1, 3–8, 10
αPV: α-parvalbumin. 1–8, 10–16, S8
*β*PV: β-parvalbumin. 4
Ca^2+^: calcium. 1–7, 9–16, S9
CaM: calmodulin. 12
CBP: Ca-binding protein. 2, 3, 12
CBPs: Ca-binding proteins. 13
K^+^: potassium. 2
MD: molecular dynamics. 1–3, 6–10, 15, S5
Mg^2+^: magnesium. 1–7, 9–16, S3
MM: scores molecular mechanics scores. 4, 7, 10, 11, S8
MM/GBSA: molecular mechanics generalized Born approximation. 1, 3–5, 14–16, S6
PV: parvalbumin. 1, 2, 12
RMSF: root mean squared fluctuations. 3, 4
SASA: solvent accessible surface area. 7
TnC: troponin C. 3, 11, 12
WT: wild-type. 1, 4–8, 10, S8

